# Long-term human organotypic brain slice cultures: a detailed protocol to provide a comprehensive framework for single-neuron and neuronal network investigations

**DOI:** 10.1101/2023.10.10.561508

**Authors:** Aniella Bak, Henner Koch, Karen M. J. van Loo, Katharina Schmied, Birgit Gittel, Yvonne Weber, Niklas Schwarz, Simone C. Tauber, Thomas V. Wuttke, Daniel Delev

## Abstract

**Background:** The investigation of the human brain at cellular and microcircuit level remains challenging due to the fragile viability of neuronal tissue, inter- and intra-variability of the samples and limited availability of human brain material.

**New method:** Here, we present an optimized work-up to use resected tissue from brain surgeries for live cell experiments in vitro.

**Comparison with existing methods:** We provide a reworked, detailed protocol of the production, culturing and viral transduction of human organotypic brain slice cultures for research purposes.

**Results:** We highlight the critical pitfalls of the culturing process of the human brain tissue and present results on viral expression, single-cell Patch-Clamp recordings, as well as multi-electrode array recordings over a prolonged period of time. Additionally, our statistics show that brain tissue from patients of any age and morbidity can be used for organotypic brain slice cultures if carefully selected.

**Conclusions:** Organotypic brain slice cultures are of great value for basic neuroscience and disease modeling over a time course of three weeks.

**Highlights:**

- Long-term human organotypic brain slice cultures are viable for 2-3 weeks and provide a framework for basic neuroscience and disease modeling
- We provide a reworked, detailed protocol for organotypic human brain slice culture production and maintenance
- We show results of long-term culturing of human organotypic brain slice cultures in terms of viral transduction, whole-cell Patch-Clamp recordings and multi-electrode array recordings
- Statistics of 16 surgeries show the correlation between the overall success rates of efficient culturing and viral transduction with the patient age and morbidity

Figure 1, Graphical abstract.
Schematic overview of preparation, maintenance and experimental workup of human organotypic brain slice cultures.
After surgical resection, human brain tissue is prepared, sliced and cultured at air-liquid-interface between human cerebrospinal fluid and defined incubator atmosphere, allowing for week-long viability. This enables extensive experimental workup such as viral transduction, single cell and multi-electrode array electrophysiological recordings.

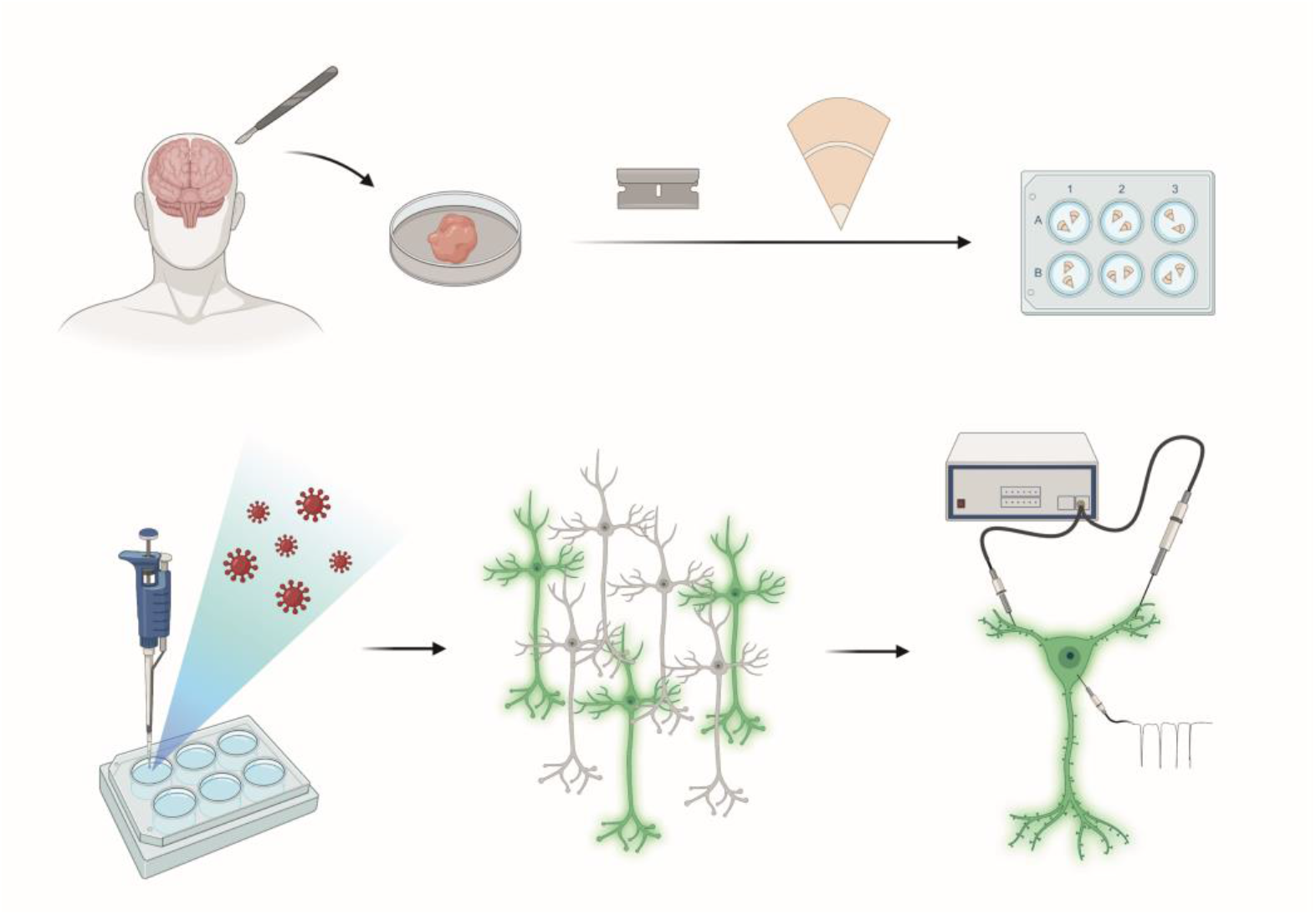

## Introduction

Keeping neuronal networks intact and functional *in vitro* over extended periods of time has been an important technical challenge in neuroscience over the last decades [1]. While culturing procedures for neonatal rodent nervous tissue have been available for a long time [2]–[4], these techniques proved to be more difficult for adult neuronal tissue [1]. Nevertheless, the culturing of brain slices has opened the avenue to study several aspects of physiology and disease *in vitro*, which was not feasible in acute slice preparations. Such so-called organotypic slices are now used for research on a wide range of neuroscientific areas, including hippocampus, cortex, spinal cord and cerebellum to disease modeling including traumatic brain injury [5], epileptogenesis [5], neurodegeneration [6], [7] and tumor biology [8].

Human brain slices from surgical tissue samples have been extensively studied *in vitro* from single cells to small networks to address neurophysiological questions [9]–[15]. Of note, most of these studies used acute slice preparations creating an investigation time window of maximum 24 - 72 hours [16]. Hence, long-term organotypic slice culturing of human brain slices is considered of great interest, as it can extend the experimental time frame of this valuable tissue considerably, opening the possibility for manipulation of human cells, for example by viral approaches. First studies have established human brain slice cultures as a matrix for investigating malignant glioma cell invasion [17] or glia cell anatomy [18]. However, successful studies of neurons and neuronal networks within slice cultures depend critically on their structural and electrophysiological integrity over the extended time *in vitro*. Eugene et al. first demonstrated preserved activity of cultured human organotypic hippocampal slices in a defined medium for up to 29 days in culture [19], which was followed by several groups using human organotypic slice cultures for electrophysiological measurements [20]–[24]. While culturing in defined medium seemed to be effective for hippocampal slice cultures, it did not yield the same results for cortical slice cultures [19]. Instead, human cerebrospinal fluid (hCSF) was demonstrated to have stimulatory effects on neuronal activity [25]–[27] and to support neuronal survival in rodent organotypic slice cultures [28]. Indeed, the use of hSCF as culture medium has a beneficial effect on the survival and activity levels of the human organotypic cortical slices [22] and also substantially increases the activity levels in the slices [21]. In first studies following this initial observation, we have started to apply human organotypic brain slice cultures towards optimization of viral transduction strategies [23], enabling us visualization and manipulation of neuronal tissue as well as single-cell and MEA investigations of human brain tissue in time and space.

Here, we describe in detail and highlight critical steps and pitfalls of the procedure of production, culturing and viral transduction of human organotypic brain slice cultures, provide a detailed guideline and present results of viral expression and electrophysiological recordings after prolonged cultivation *in vitro*.

## Materials & Methods

### Preparations one day prior to surgery

For the preparation of cortical and/or hippocampal slices of the human brain, a choline containing slicing solution is used, as choline is supposed to diminish slicing stress on neurons by replacing sodium and thus, silencing the neuronal tissue and decreasing excitotoxicity [29] (Suppl. A, note 1). The human brain slicing solution (s-aCSF) contains 110 mM choline chloride, 26 mM NaHCO_3_, 1.25 mM Na_2_HPO_4_, 11.6 mM sodium ascorbate, 3.1 mM sodium pyruvate, 7 mM MgCl_2_, 0.5 mM CaCl_2_, 2.5 mM KCl, 10 mM glucose and 1% penicillin/streptomycin/amphotericin B dissolved in Millipore water. Subsequently, the solution is carbogenated (95% oxygen, 5% CO_2_) for at least 20 minutes before setting the pH to 7.4 with HCl and NaOH respectively. Finally, the s-aCSF is sterile filtered (filter pore size 0.2 µm). Two liters of s-aCSF should always be prepared fresh either on the day of surgery or the day before at maximum to prevent the choline from precipitating and can be stored at either 4°C or -20°C.

Solidified agarose is prepared at a concentration of 4% according to standardized protocol the day before surgery to stabilize brain tissue during the slicing process.

Regular artificial cerebrospinal fluid (aCSF) is used to store human brain slices in-between slicing and culturing, as well as for electrophysiological recordings. The aCSF contains 125 mM NaCl, 25 mM NaHCO_3_, 2.5 mM KCl. 1.25 mM NaH_2_PO_4_, 1 mM MgCl_2_ * 6 H_2_O, 2 mM CaCl_2_, 25 mM glucose and 1% penicillin/streptomycin/amphotericin B dissolved in Millipore water. Penicillin/streptomycin/amphotericin B are omitted from aCSF used for electrophysiological recordings. The solution is carbogenated (95% oxygen, 5% CO_2_) for at least 20 minutes before setting the pH to 7.4 with HCl and NaOH, respectively. Finally, the aCSF is sterile filtered and stored at 4°C until further use. aCSF can be stored up to several weeks as long as pH and osmolarity (310 - 320 mOsm/l) stay in range.

All tools needed for tissue preparation, culturing and/or slicing are sterilized beforehand by means of autoclaving or sterilizing oven (glass/metal tools) or UV light.

### Culture preparation on the day of surgery

All following steps are carried out under a thoroughly cleaned sterile hood (Suppl. A, note 2). Human brain slices are pre-incubated in medium highly supplemented with HEPES before being placed in long-term culture to provide additional buffering capacity and allow for the stabilization of a physiological pH directly after slicing. For the intermediate step HEPES medium (ISHM), 23.5 ml DMEM/F-12 and 23.5 ml Neurobasal medium are mixed. Subsequently, 1 ml B27 supplement, 0.5 ml N2 supplement, 0.5 ml GlutaMAX, 0.5 ml penicillin/streptomycin and 0.5 ml non-essential amino acids are added to the medium to yield 50 ml of ISHM. To establish buffering capacity, 20 mM of HEPES are added to the medium and stirred in for at least 20 minutes. Finally, the medium is adjusted to a pH of 7.4 with HCl and NaOH respectively, sterile filtered and stored at 4°C until further use. The ISHM should always be prepared fresh on the day of surgery.

Human cerebrospinal fluid (hCSF) used for culturing is mostly collected from lumbar punctures for pressure relief of patients with idiopathic intracranial hypertension or for diagnostic workup of normal pressure hydrocephalus (NPH). Written and informed consent is collected for every patient in accordance with the approval of the local ethics committee (EK064/20). After collection, hCSF should directly be processed (see Note 3). For use as human brain slice culturing medium, hCSF is centrifuged at 4°C and 2000 g for 10 minutes and sterile filtered subsequently (0.2 µm pore size). Osmolarity for every batch of hCSF is controlled and should be at 280 mOsm/l with a deviation range of +/- 20 mOsm/l [30]. Readily processed hCSF is then stored in aliquots of 10 ml at -80°C until further use.

For culture preparation, the number of planned wells on 6-well plates are filled with 1.5 ml freshly prepared ISHM medium each. Subsequently, cell culture inserts with hydrophilic semi-permeable membranes (0.4 µm pore size, 30 mm diameter) are carefully placed on the medium surface without trapping air bubbles underneath. The plates are placed in the incubator (37°C, 5% CO_2_ and 100% humidity) to equilibrate temperature and oxygen/CO_2_ within the medium. For long term culturing, processed hCSF is thawed in a water bath and the equivalent number of 6-well plates are filled with 1.5 ml hCSF per well as done for the ISHM.

### Slicing preparation on the day of surgery

All following steps are carried out on a clean working bench and should be performed as sterile as possible. All tools and required items were sterilized by heat or UV light; vibratome and tabletops are wiped with 70 % ethanol (Suppl. A, note 2). s-aCSF is thawed at 4° C if frozen beforehand. For use with tissue, s-aCSF should be ice-cold, but not frozen or “slush” anymore, as ice crystals could damage the tissue. The working place is lined with absorbent sheets and on top, vibratome, binocular, preparation dishes, preparation tools, glue, parafilm and glass pipettes are set up. The vibratome is equipped with a double-edged razor blade and z-axis deflection is controlled with VibroCheck. The slicing chamber of the vibratome is filled with ice-cold s-aCSF and additionally ice cubes are added for prolonged cooling (Suppl. A, note 4). Preparation dishes and bottles are filled with ice-cold s-aCSF and ice cubes as well. Ten falcons are filled with s-aCSF for rinsing and kept on ice. A slice keeper is set up on a controlled heating plate and filled with aCSF, heated to a stable temperature of 30 - 33°C. A new, sterile tubing system is installed from the main carbogen gas tap to carbogenate the vibratome slicing chamber as well as every preparation dish and slice keeper (Suppl. A, note 5). For this cause, three-way valves and clinical infusion sets (or other sterile tubing - must be without air trap) is used with medium cannulas installed at the ends, which are then placed into the buffer for regulated airflow. Subsequently, carbogenation of slicing chamber, preparation dishes and slice keeper is started. Agarose blocks are cut into small square cubes, which should be small enough to fit on the magnetic disc of the vibratome, but big enough to hold tissue blocks.

### Surgical procedure

Surgical cases are carefully and properly selected by the neurosurgeon in charge based on the type and localization of the lesion and the surgical strategy for resection (Suppl. A, note 6). Generally, surgical cases for removal of pathologies in the vicinity of eloquent areas, large vascular structures or bony parts like temporal base are not suitable. Frequently, it may be required to resect superficial cortical tissue during the surgical approach to the lesion. Such tissue often is removed and destroyed by use of an ultrasonic surgical aspirator. However, alternatively dissection and en-bloc resection may be possible such that these samples can be used for human organotypic brain slice culturing. Informed and written patient consent for tissue donation and approval of the use of human tissue for scientific purposes and for all related experimental procedures by respective institutional ethics boards was collected before commencement of the study (EK067/20).

In case tissue from different locations has to be removed due to the surgical strategy/approach, collecting samples that can be quickly dissected and with minimal or without use of bipolar cauterization (i.e. fully exposed tissue of the cortical convexity rather than aspects of the cortex adjacent to the skull base that are harder to reach) is prioritized. Trepanation and opening of the dura are performed according to standard neurosurgical practice. From there on, there are three main possibilities to obtain the brain tissue (Suppl. A, note 7). Cortical tissue can be gathered either as a small tissue block or as en-block resection of larger tissue parts. The third option refers to the preparation of a hippocampus.

In cases of small tissue block resections (Fig. 2A-D), coagulation with the bipolar forceps should be completely avoided. Under the microscope, the pia is cut with blade #11 in rectangular form measuring 0.5 – 1.0 cm length of each side. The macro-dissector is used to dissect the block of tissue going approximately 1-2 cm deep in the tissue. For larger en-block resections, a complete avoiding of coagulation with the bipolar forceps is difficult. However, only the edges of the resection should be coagulated and resected. To dissect the tissue, ultrasonic aspiration is used and coagulation should be performed only to the sulcular vessels before cutting them. The obtained tissue blocks are immediately transferred to ice-cold, carbogenated s-aCSF. The tissue block should not be squished or exhibited to any kind of mechanical stress during transfer (Suppl. A, note 8).

**Figure 2. A-D:**
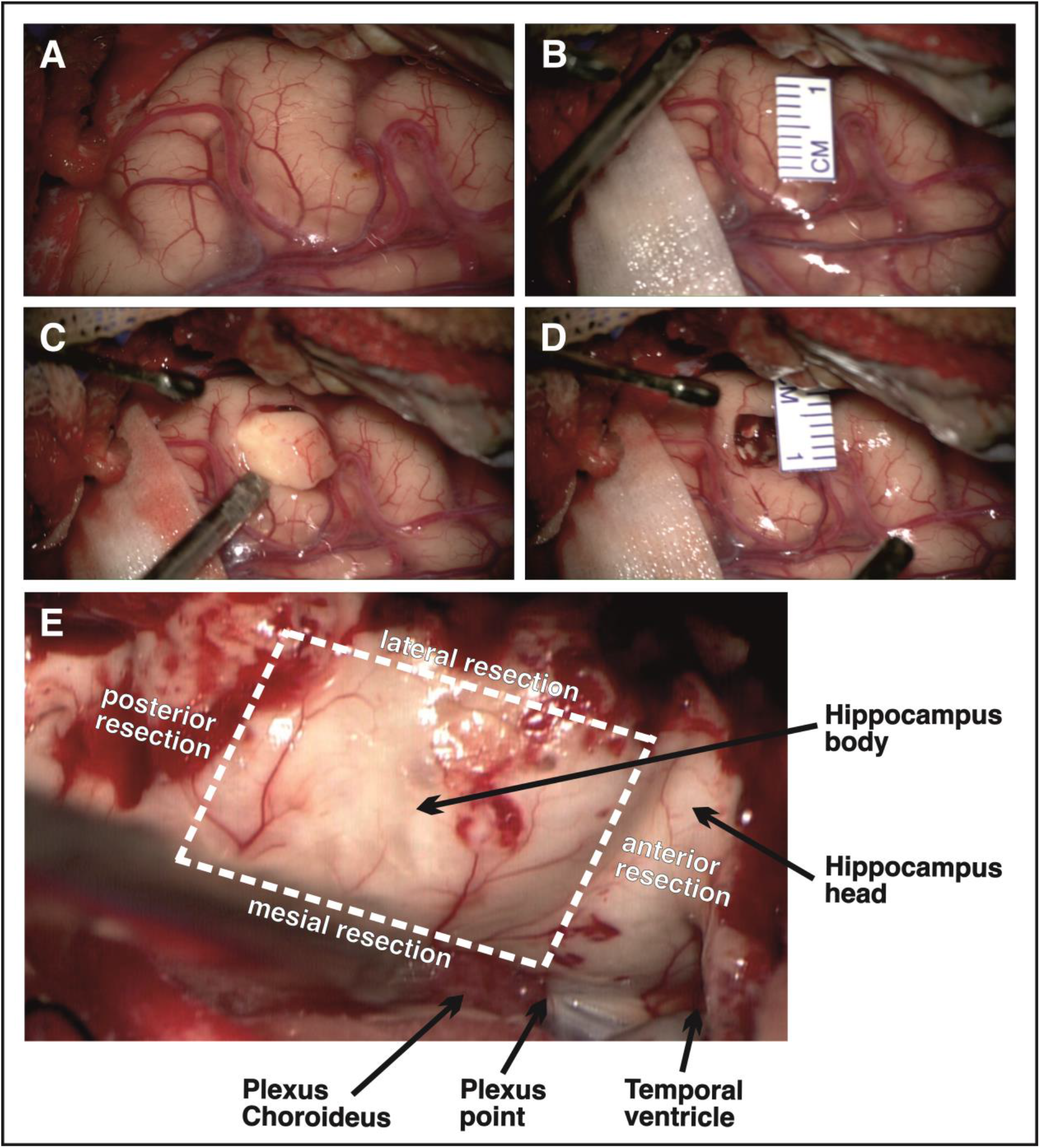
The resection of the cortex is presented. A) A part of the cortex was selected, which is not marked by vessels. Sulci are avoided due to difficulties concerning the preparation of the pia. B) Measurement of approximately 1 cm of cortex. C) Resection is performed without any electric or thermal coagulation. D) Cavity after the removal of the tissue. E) **En-block removal of the hippocampus.** Important anatomic landmarks (Plexus choroideus, Plexus point, temporal ventricle) should be identified. The border between hippocampus head and body should be identified as well. The dashed lines show the resection borders: lateral = Gyrus parahippocampalis, mesial = Fimbria hippocampi, anterior = Hippocampus head, posterior = Hippocampus tail.

For hippocampal resection (Fig. 2E), the temporal horn of the ventricle is widely opened to give the opportunity to identify amygdaloid body, hippocampus head, choroidal point and fimbria of the hippocampus. The amygdala is resected and the hippocampus head is removed just before the choroidal point with the ultrasonic aspirator. At this point, the hippocampal fissure is identified. The lateral disconnection is performed by resecting the parahippocampal gyrus until the temporal basal pia is reached. The dorsal disconnection is executed approximately 2.5 cm in the longitudinal projection of the hippocampus. The hippocampal fissure is identified and preserved. Finally, the mesial disconnection is performed. For this, the fimbria was elevated with the dissector and the hippocampus body is carefully pealed from mesial to lateral in a subpial manner until the apex of the hippocampal fissure is reached. The vessels in the hippocampal fissure are coagulated proximal to the hippocampus and cut with micro-scissors.

### Transfer of tissue from surgery to laboratory

For transportation of surgically removed brain tissue from the operating room to the laboratory, autoclaved bottles with ice-cold s-aCSF are placed in a Styrofoam container filled with ice to keep the solution cold. The transportation setup should arrive at the operating room before the resection is fully completed. Carbogenation of the s-aCSF buffer is started at least 15 min prior to final resection. As soon as the tissue block is completely resected from the brain, it is directly placed into ice-cold, carbogenated s-aCSF. The tissue should not be left without oxygen and/or fluid for more than a few seconds. A blunt object is used to gently move the tissue into the fluid. The tissue should not be pushed, squeezed or exposed to any other mechanical stress. The bottle with s-aCSF and the tissue block is placed back into the styrofoam container with ice and moved back to the laboratory while carbogenating constantly.

### Tissue dissection and block preparation

During preparation, ice cubes are constantly replaced to keep the slicing solution cold. After arriving at the working bench, the brain tissue is gently moved from its bottle into the first falcon with ice-cold s-aCSF. The falcon is inverted carefully several times to rinse the tissue and wash off any blood or other residue. This step is repeated with the remaining nine falcons. Finally, the rinsed tissue block is moved to a dissection dish filled with ice-cold, carbogenated s-aCSF. Depending on the size of the tissue block, it can be directly dissected into smaller blocks using a scalpel or razor blade to ensure proper oxygenation. If visible, this is done best by cutting perpendicular to the tissue surface along the Sulci and/or crosswise through the gyrus (Fig. 3 A1 - A2).

**Figure 3.**
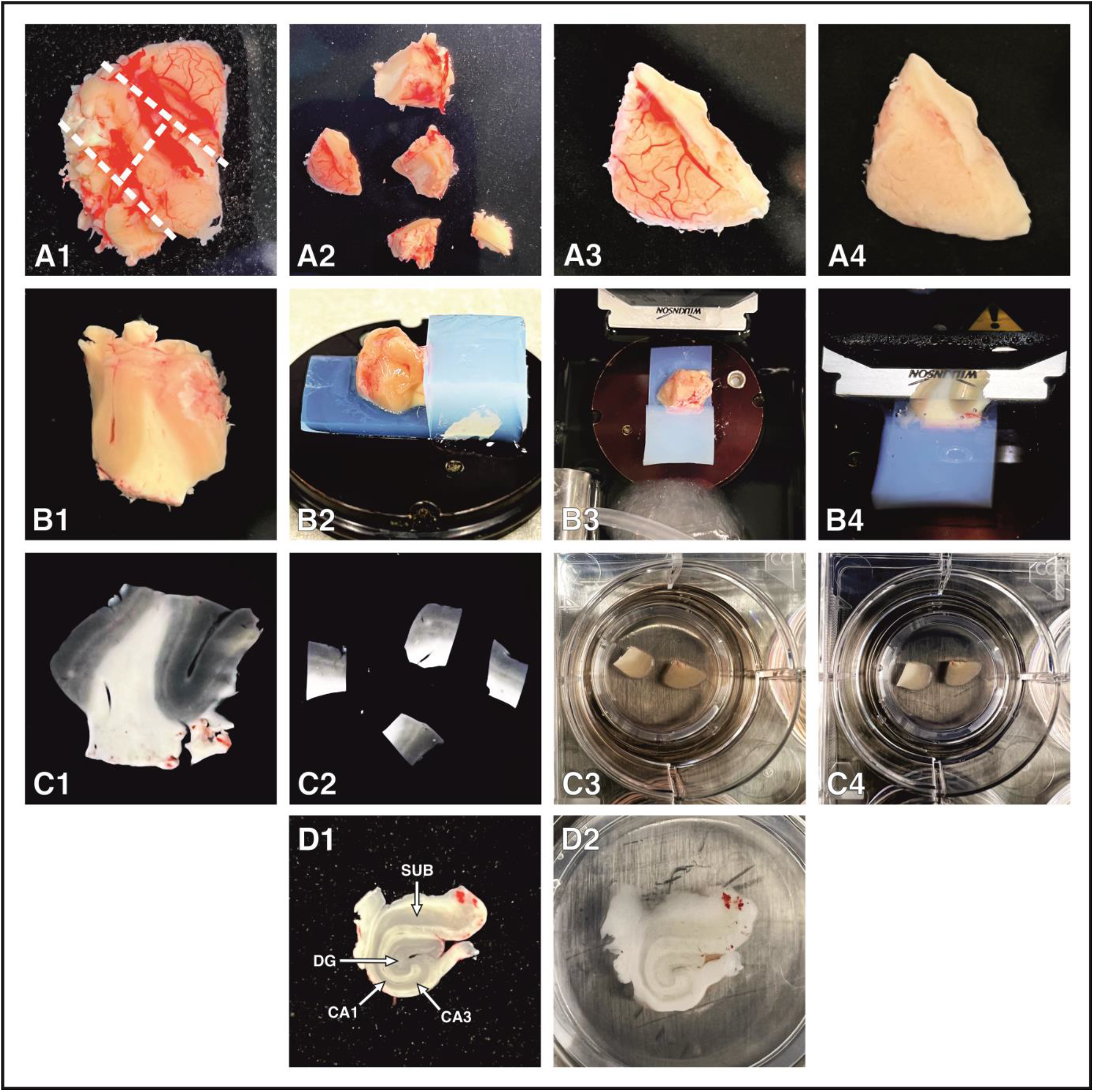
Step-by-step visual representation of human brain tissue preparation, slicing and culturing. A1) Brain tissue block as resected by the surgeon. Separation into smaller blocks along the dotted lines. A2) Resulting brain tissue blocks with correct orientation after perpendicular separation. A3) Brain tissue block with clearly visible meninges and vessels on cortical surface. A4) After removal of meninges and vessels, the cortical surface should appear smooth and unichrome. B1) Flat and correctly oriented contact surface of brain tissue block before slicing. B2) Positioning of stabilizing agarose blocks and tissue thereon. B3) Orientation of vibratome metal plate within the slicing chamber with brain tissue facing the blade. B4) Brain slice during slicing process. C1) Uncut brain slice directly after slicing with white matter and imperfections still attached. Cortical layer 4 clearly visible as light streak traversing the cortex. C2) Finished brain slices after cutting and trimming to achieve desired size and condition. C3) Intermediate step HEPES medium with two brain slices placed on top of semipermeable membrane insert. C4) Culturing insert with brain slices after moving from intermediate step medium to hCSF for long term culturing. D1) Acute brain slice from human hippocampus directly after slicing and encompassing CA1 – CA3. B) Hippocampal brain slice 7 days *in vitro*. Note that no obvious structural changes are visible.

Leptomeninges (Fig. 3 A3) are removed as completely as possible by grabbing them with forceps at the cutting edges and gently pulling. Cortex is held down carefully with a blunt spatula while doing so to not damage the tissue parenchyma along with the arachnoidea/Pia mater. If the Pia mater won’t disconnect from the tissue underneath, it can be cut in small pieces with microdissection scissors. The final cortex surface should be looking smooth and homogenous, without Pia mater or any vessels left (Fig. 3 A4). For slicing of cortical resections, tissue blocks of max. 0.75 x 0.75 x 0.75 cm or smaller are most suitable. To obtain a clear orientation of the brain slices later on, tissue blocks are cut straight from the cortex surface towards the white matter perpendicular to gyrus anatomy/direction. The result should be a straight contact surface with cortical layer 4 clearly visible as light streak traversing the gray matter and gray matter being distinctly separated from white matter (Fig. 3 B1). For optimal hippocampal slices, the hippocampus should be resected en block. The Hippocampus is subsequently glued on an agar block to achieve stability and cut in 250-300 µm thick slices. The slices should contain dentate gyrus (DG), CA1 – CA3 and subiculum (Fig. 3 D1).

### Slicing procedure

During slicing, ice cubes are constantly replaced to keep the slicing solution cold. To support the brain tissue block while slicing, a thin agarose block is superglued flat on the magnetic plate of the vibratome. Afterwards, another thicker agarose block is glued upright to the edge of the thin agarose block. The prepared brain tissue block is placed on the thin agarose block with the oriented flat contacting surface facing downwards. Subsequently, the brain tissue is pushed back against the thick agarose block gently with its straightest side (Fig. 3 B2). With a bent cannula, a minimal amount of glue is taken up and placed on the thin agarose block, very carefully pushing it against the edges of the tissue block to secure it into place. The magnetic plate with the mounted brain tissue is placed into the slicing chamber of the vibratome, filled with ice-cold, carbogenated s-aCSF, with the tissue facing the blade and the thick agarose block facing away from the blade (Fig. 3 B3). Slicing of brain tissue is started from top to bottom (Fig. 3 B4) using chosen parameters (example parameters: 250 µm slice thickness and 0.1 mm/s speed). As soon as the slice dissociates from the block (Suppl. A, note 9), it is taken up with a wide mouth glass pipette and moved to a fresh preparation dish. Do not exhibit any kind of mechanical stress on the slices by pipette blowing or handling them with other objects. In the preparation dish, slices are cut (Fig. 3 C1) into desired shape and size using a scalpel or razor blade (Fig. 3 C2). If the cortex seems to be damaged in certain locations, these parts are removed with a straight cut. Until enough slices are collected to start the culturing, the readily made slices are put into the slice keeper filled with pre-warmed and oxygenated aCSF.

### Culturing procedure

All following steps are carried out under a thoroughly cleaned sterile hood. Slices are transferred onto inserts in 6-well plates for pre-incubation with ISHM. Brain slices should be positioned flat and completely spread out on the membrane. If this is not the case, slices should not be moved by pushing with any object, but rather carefully by taking it up again with the pipette and some fluid to reposition it. Per insert, 2 - 3 brain slices (depending on size) can be placed on the membrane without touching each other (Fig. 3 C3). The excess s-aCSF on the membrane insert is then carefully removed using a 200 µl pipette, which leads to adhesion of the slices to the membrane, creating a liquid-air interface. As soon as the positioning of slices in the 6-well plate is finished, the plates are placed into the incubator. Brain slice cultures are then incubated on ISHM for at least one hour. After ISHM incubation, inserts holding the brain slices are carefully taken out of their well and most ISHM is removed from their top and bottom side. Inserts are slowly placed into the wells with pre-incubated hCSF for long-term-culturing (Fig. 3 C4). Medium change is performed every 2-3 days by removing 750 µl of hCSF from the wells and replacing it with 750 µl of pre-warmed fresh hCSF. The condition of the slices and transparency of the medium should be monitored continuously during the culturing to recognize possible degradation or infections (Suppl. A, notes 10 and 11).

### Viral transduction

Viruses used for human organotypic brain cultures do not have to fulfill any generalizable requirements. Both commercially available and self-produced viruses (concentrated or the supernatant of the virus production culture) can be used. Transduction efficiencies, expression levels and time course of expression will depend on serotypes/capsids and tropism, titers, promoters, construct engineering and on whether self-complementary vectors are being used or not. In our experience, adeno-associated viruses (AAVs) and lentiviruses have yielded robust results so far, however, herpes simplex viruses (HSV) were also shown to be effective [20]. Viruses should be distributed in aliquots after the first thawing to avoid unnecessary freeze-thaw cycles.

There are two possible methods for virus application (Suppl. A, note 12): The first option is the application of the virus (1-2 µl) to the surface of the slice with a pipette tip. In this case, the pipette tip must not touch the tissue, but only place the droplet on the brain slice at the front of the tip resulting in a relatively wide transduction area.

The second option allows a much more precise application of the virus by using a glass pipette. For better visualization, the virus can be mixed 10:1 virus/dye ratio with FastGreen dye to monitor the spread in the tissue. The glass pipette tip is guided into the tissue by manipulators and the application is achieved by a short pulse of compressed air. The preferable tool for this approach is a picrospritzer, where the pressure and duration of the pulse can be adjusted.

## Results

### Viral expression

Viral transduction of organotypic human brain slice cultures can serve as a helpful readout to determine the viability and integrity of the endogenous neuronal network, but can also be used as a versatile tool to introduce molecules of interest into target cells. Possible molecules to (over)express in human brain tissue include, but are not limited to fluorescent reporters, calcium indicators, opsins and short-hairpin RNAs, and can be expressed under ubiquitous, neuron-specific or even cell type-specific promoters. Simple fluorescent markers such as GFP expressed under a neuron-specific promoter (e.g. the human Synapsin-1 (hSyn) promoter) will result in wide-spread expression in cortex (Fig. 4A) as well as hippocampus (Fig. 4B), and can be routinely used to observe the structural integrity of the neuronal network. In stable and viable organotypic brain slice cultures, neurons should always appear fully intact with most structures such as the apical dendrites clearly showing fluorescence activity (Fig. 4C). If neurons appear fragmented and dendrites represent in individual, disconnected parts (Fig. 4D), slice cultures are degrading and as a result, neurons are disintegrating. The rise of AAV-mediated expression can vary and, just like general culture quality, depends on parameters such as tissue quality, preparation quality, patient age and previous radiation and/or pharmaceutical treatment among others. Nevertheless, expression can be usually expected between 3-5 days post-transduction.

**Figure 4.**
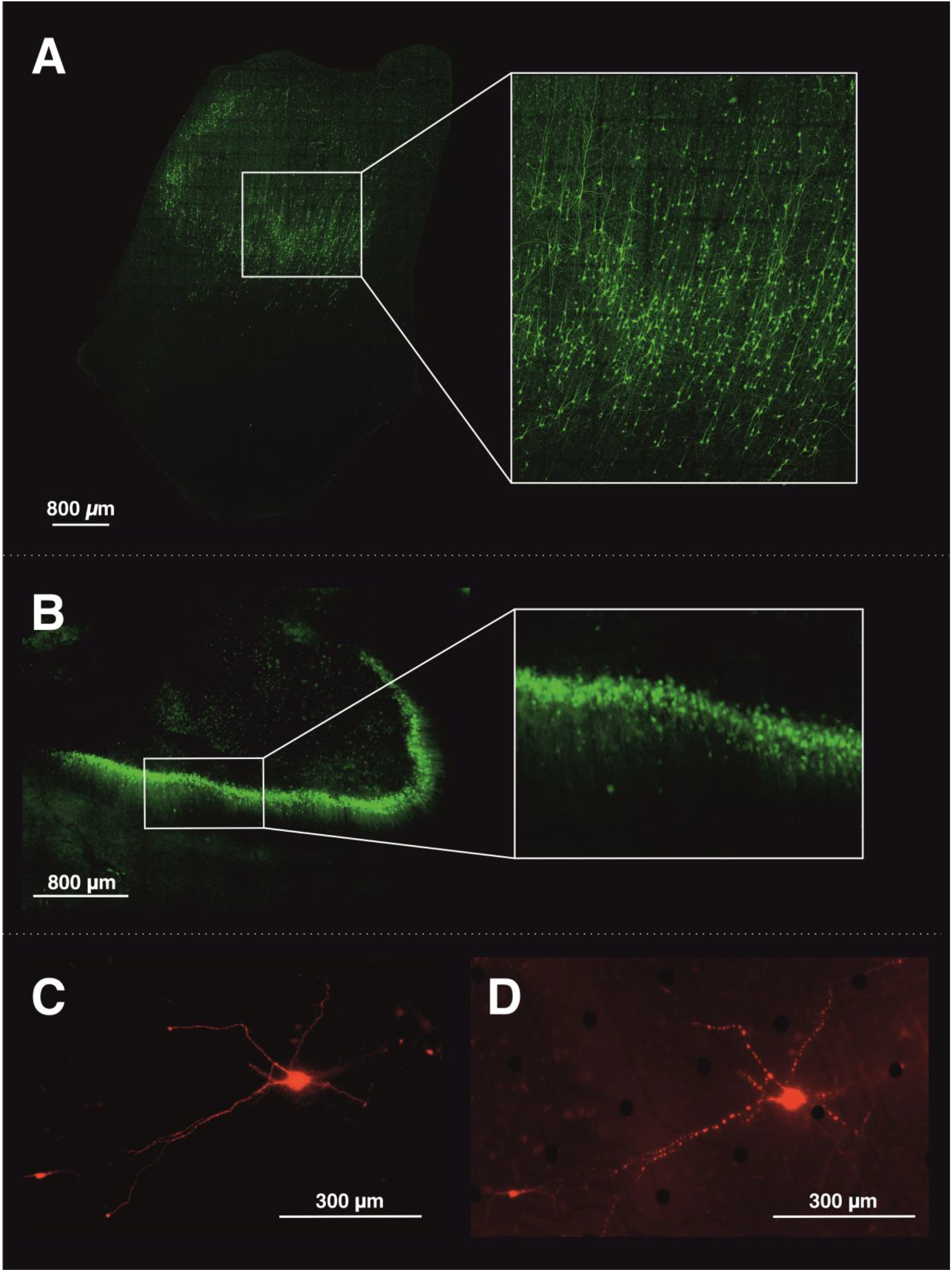
Viral expression of fluorescent reporters. A) Human cortical brain slice transduced with retrograde AAV encoding fluorescent GFP under hSyn promoter (transduced at 1 DIV, imaged at 7 DIV). Wide-spread GFP expression among all cortical layers of the slice. B) Human hippocampal brain slice, transduced with retrograde AAV encoding fluorescent GFP under the hSyn promoter (transduced at 1 DIV, imaged at 15 DIV). C) A human cortical neuron expressing mCherry under under the hSyn promoter; example for intact, viable cell. D) A human cortical neuron expressing mCherry under the hSyn promoter; example for fragmented, disintegrating cell.

### Electrophysiological recordings

Our model of human brain slice cultures provides a good opportunity for any electrophysiological investigation since its viability can last for up to three weeks. Therefore, electrophysiological activity is an important parameter to determine the viability of organotypic brain slice cultures as well as a valuable readout for corresponding experiments. There are several approaches to assess the electrophysiological activity of brain slices on multiple levels, from the cellular to the whole-slice network level. Here, we were able to implement and show recordings from two commonly used electrophysiological techniques: intracellular Patch-Clamp recordings and multi-electrode array (MEA) recordings.

#### Cellular parameters

*Intracellular Patch-Clamp recordings.* Evaluation of basic cellular properties such as membrane potential, input resistance, firing behavior and spontaneous activity can be performed using intracellular Patch-Clamp recordings. They deliver information about the general condition and viability of single cells and, additionally, offer the possibility of a variety of chemical and electrical modulation to further assess neuronal properties. Furthermore, the visual Patch-Clamp technique also allows for a first impression of the cellular morphology and a rough estimate of the number of viable cells within the slice. Patch-Clamp recordings can also be combined with viral transduction of a fluorescent protein a few days prior to the recordings to enable easier visualization of the neurons, especially those located in the depth of the slice. Imaging of single cells however also is possible if either a fluorescent dye or another compound such as biocytin is added to the intracellular solution used for the recording to stain the patched cell afterwards. This allows for a precise visualization of single neurons, complete neuronal reconstruction (Fig. 5A) as well as their exact morphological characterization and comparison with firing properties. If the brain slice cultures are viable, neurons will sustain physiological firing properties over the cultivation period of several weeks (Fig. 5 C-F) and also exhibit spontaneous network activity (Fig. 5E).

**Figure 5.**
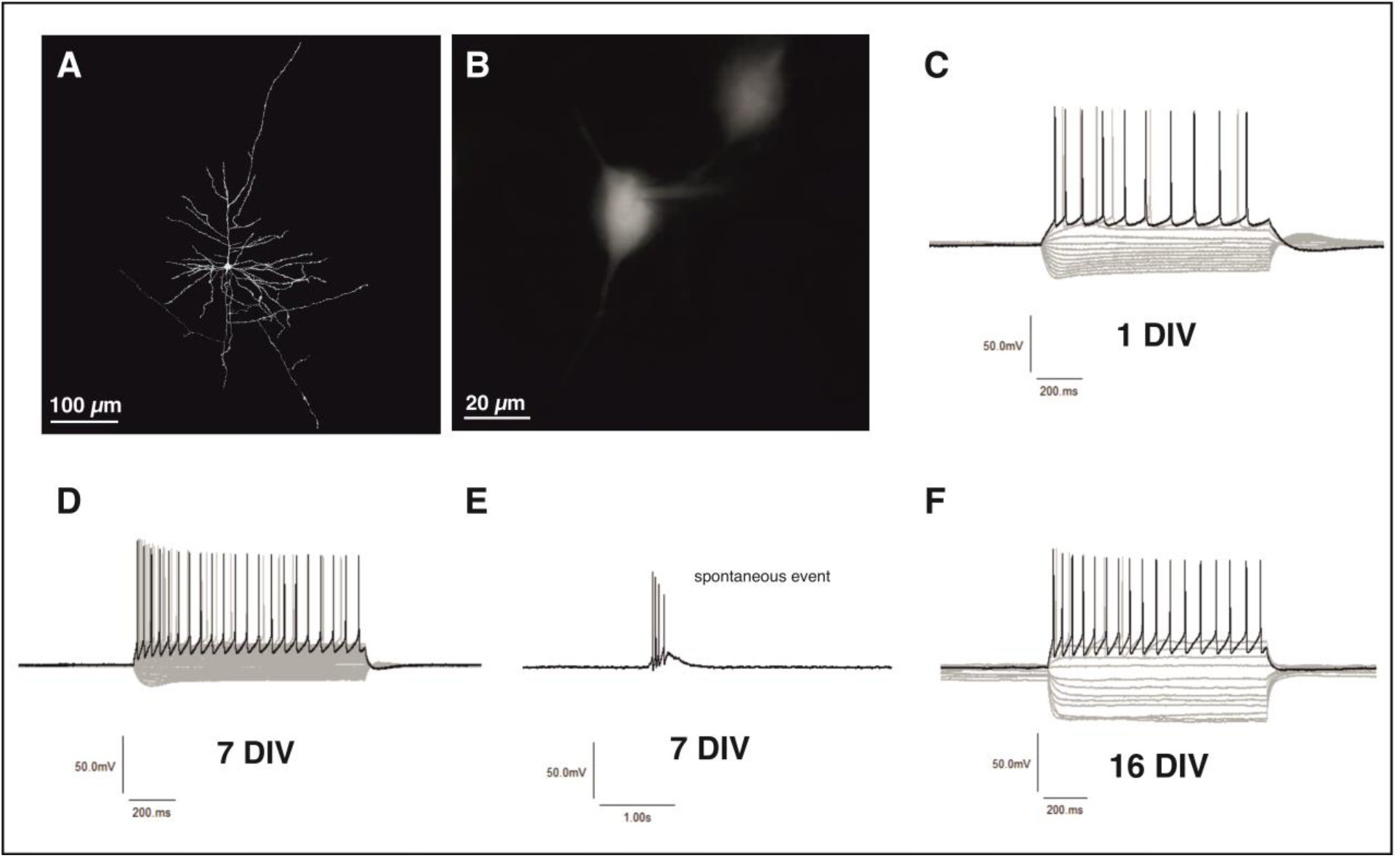
Physiological firing properties of human neurons in organotypic brain slice cultures. A) A layer 2/3 pyramidal neuron was filled with biocytin during a whole-cell Patch-Clamp recording and subsequently stained with Streptavidin-Cy3. Neuronal reconstruction of the recorded cell. B) Close-up of human pyramidal cell during whole-cell Patch-Clamp. C) Example traces from whole-cell Patch-Clamp recordings of a pyramidal neuron portrayed at 1 DIV. Action potentials fired after stimulation pulses, beginning at -200 pA with 25 pA steps. D) Example traces from whole-cell Patch-Clamp recordings of a pyramidal neuron portrayed at 7 DIV. E) Spontaneous events recorded without stimulation in current-clamp mode. F) Example traces from whole-cell Patch-Clamp recordings of a pyramidal neuron portrayed at 16 DIV.

#### Whole-slice dynamics

*Multi-electrode array (MEA).* To better determine network dynamics in the slice cultures, the MEA system can be used. The spacing of the electrodes can be varied and fitted to the slice culture size. The human cortical slices cultures can be well covered with electrodes (Fig. 6A), e.g. 16 x 16 electrodes = 256 electrodes (size 30 μm, spacing of 200 μm), covering a recording area of ∼3.2 × 3.2 mm^2^ (e.g. USB-MEA 256-System, Multi-Channel Systems MCS GmbH). The MEA technique enables recordings of the local field potential (LFP) and action potential (AP) activity along large parts or even the entirety of the slice surface (Fig. 6B), as well as analysis of onset and termination points, propagation properties and amplitudes of an event (Fig. 6C-E) and is a powerful tool for the investigation of complex neuronal network activity.

**Figure 6.**
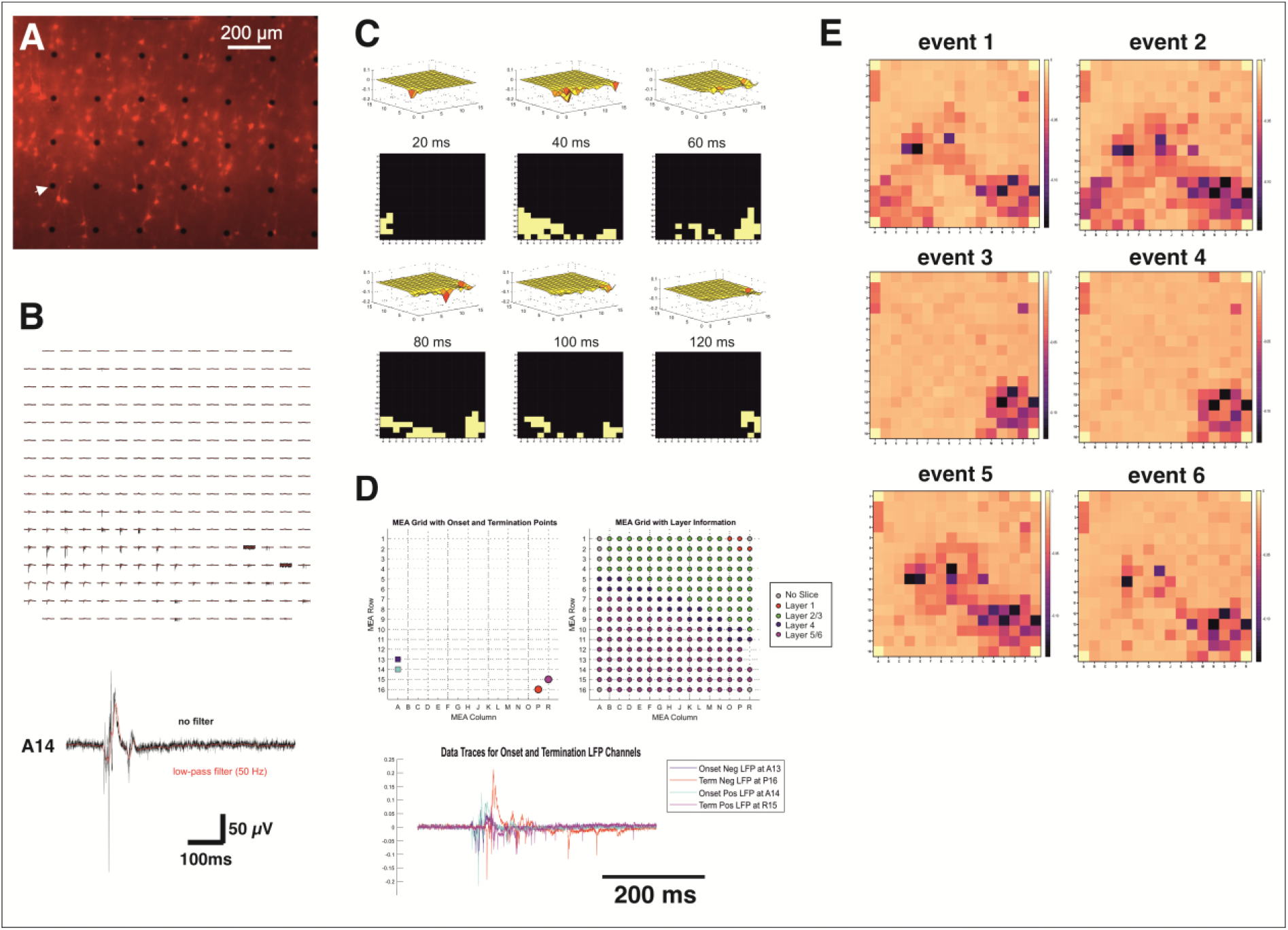
Network activity recorded using the MEA system. (A) Example of a live-imaging picture of mCherry labeled neurons in human organotypic slice culture in relation to the recording electrodes of the MEA (black dots, see white arrow head). (B) Example of synchronous discharges over a large part of the MEA grid (top) and an example of one channel of a synchronous event (red low-pass filter and black without a filter), note the slow LFP component and the fast AP component. (C) Example of the propagation of the LFP signal over the MEA grid. (D) Representation of the onset and termination of the LFP signal on the MEA grid and correlation to the layers of the cortex. (E) Displayed are the negative amplitudes of the LFP signal in subsequent events, note the similarities and variability in the spatial distribution of the LFP signals.

### Overall success rate of culturing organotypic human brain slices

The cultivation of human organotypic brain slices can be a challenging method and requires an elaborate process of establishment, however, once a stable standard operation procedure is achieved, it can serve as a method holding great potential for many different applications. To demonstrate the potential output and success rate of this method, we analyzed the outcome of the past 16 surgeries (7 epilepsy, 9 tumor resection surgeries) of which we attempted to culture human organotypic brain slices. This data set covers an age range from 22 – 74 years, a gender distribution of 69% male and 31% female patients (Fig. 7A) as well as tissue resected from frontal, temporal, parietal and occipital lobes, however, most samples were obtained from temporal lobe resections (69%). We defined general success of cultures in three categories, namely *i)* successful electrophysiological recordings between 3-7 days *in vitro*, *ii)* successful viral expression, and *iii)* an overall stability of the cultures (meaning electrophysiological recordings and/or viral expression still stable) beyond 7 days *in vitro*. No clear correlation between culturing success and gender or resection area could be found, however, we found a slight decrease in the overall success rate in tissue obtained from patients with advanced age (60 – 80 years) (Fig. 7B). The majority of cases were evenly distributed in terms of age and morbidity (Fig. 7C). Over a course of 16 surgeries, electrophysiological recordings during the first week of culture were successful in 13 of 16 cases (success rates 85% for epilepsy, 78% for tumor surgeries), viral expression was successful in 13 of 16 cases (success rates 100% for epilepsy, 67% for tumor surgeries) and long-term stability over 7 days was achieved in 11 of 16 cases (success rates 57% for epilepsy, 78% for tumor surgeries), therefore suggesting no clear correlation between overall success rate and type of resective surgery (Fig. 7D).

**Figure 7.**
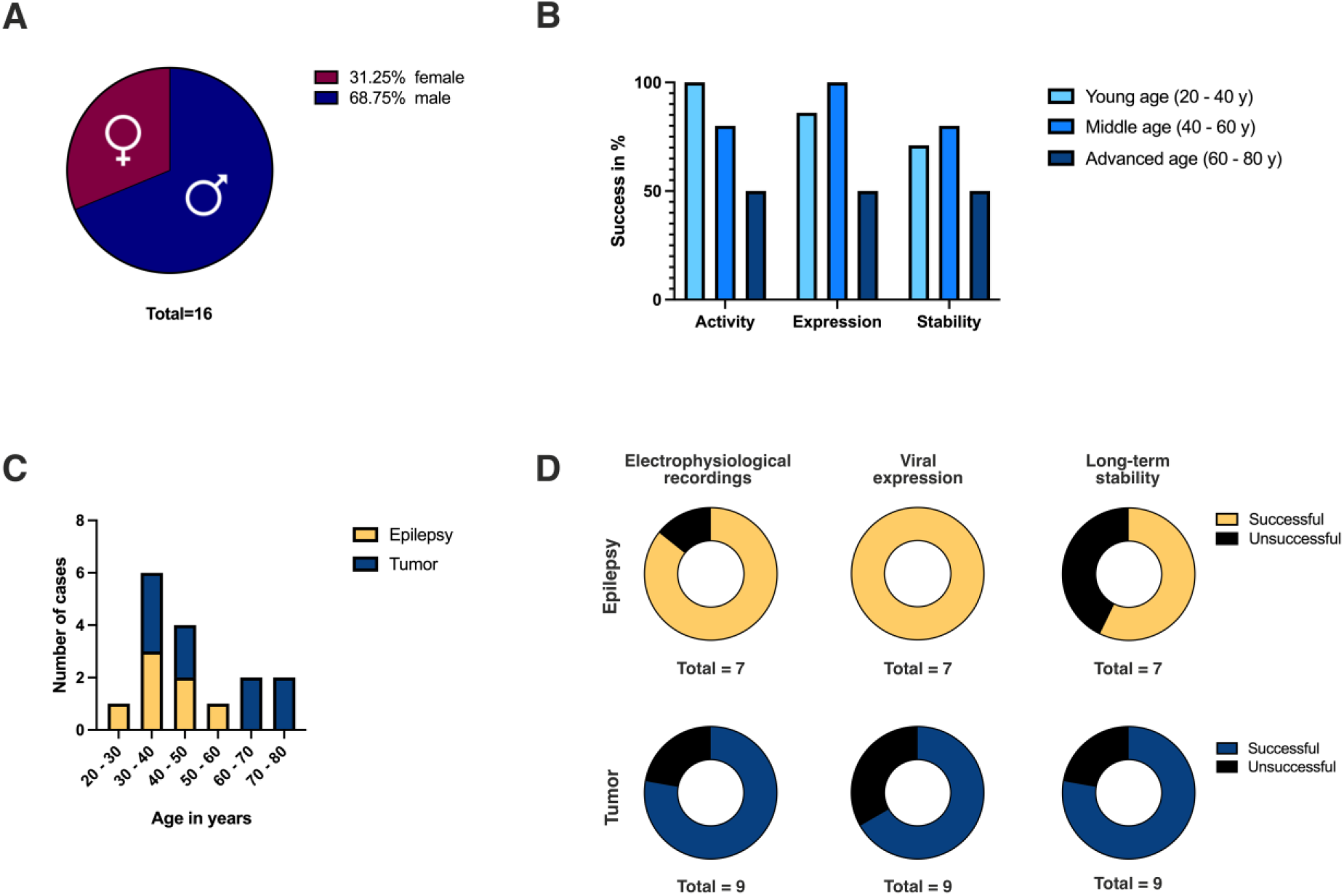
Statistical analysis of organotypic human brain slice cultures over 16 surgeries. (A) Data encompasses 16 patients, 5 female and 11 male. (B) Success rates of electrophysiological recordings, viral expression and long-term stability shown for groups of young age (20-40 years), middle age (40-60 years) and advanced age (60-80 years). No clear correlation was found except for a slightly decreased overall success rate in the advanced age group. (C) Distribution of tumor and epilepsy cases over age groups. (D) Individual success rates for electrophysiological recordings, viral transduction and long-term stability depicted for tumor and epilepsy cases.

## Discussion

The understanding of both physiological and pathological activity arising from human neuronal tissue has become a quest of increasing importance in the field of neuroscience. Intact human brain tissue allows the investigation of specific properties of the human brain from the single cell to small network level [31]–[35]. Measurements of electrophysiological activity in an acute experimental setting have been the gold standard for many years. However, there is an increasing demand for an experimental framework allowing spatio-temporal electrophysiological investigations combined with modulatory approaches, as many pathophysiological processes develop over time and show complex spatial relationships.

Following this line, organotypic slice cultures have become crucial for the investigation of single-cell and neuronal network properties of brain tissue [20], [36], [37]. Over the last few years, significant improvements have been made in establishing, maintaining, and exploiting organotypic slice cultures from rodents. However, long-term electrophysiological investigations of human brain tissue have turned out to be more challenging, since the preservation of intact neuronal activity over several days to weeks requires an extensive and elaborative work-up [23].

Here, we provide a reworked and detailed step-by-step protocol and report results of the utilization of long-term human brain slice cultures. Using access brain tissue, obtained during a surgical procedure, we were able to establish a human brain slice culture setting that can keep the neuronal activity intact for up to 3 weeks [22]. Our experimental model enables a successful viral transduction in 81% of the slices. More importantly, we are able to measure electrophysiological activity and perform single-cell and MEA recordings in approximately 80% of the slices, and even keep the neuronal activity stable in 68% of the slices for a period longer than 1 week (Fig. 7).

Although human CSF has been proven to be a valuable and well-functioning cultivation medium for organotypic human brain slice cultures [22], [28], it is a scarce resource and often not easy to obtain in sufficient quantities. Other methods have been described using either a defined cell culture medium [38] or an elaborate, defined medium mimicking the human CSF [19] to explore the possibility of culturing human organotypic brain slice cultures for a longer period of time.

### Human brain tissue availability and differences

Although the proper selection of the cases, from whom human brain tissue was obtained, is critical for the success rates of the culturing process, we could show that access tissue from different types of neurosurgical resection can be used (Fig. 7). Previous assumptions that only tissue from non-lesional cases (i.e. patients with drug-resistant epilepsy) is suitable for efficient long-term culturing are rather incorrect, since we were able to culture access tissue from patients with brain tumors as well. Therefore, our data suggest that the quality properties of the access tissue are of paramount importance for the successful culturing process rather than the type of morbidity itself. Interestingly, the success rates of electrophysiological recordings and viral expression in tissue gathered from epilepsy resections were indeed higher compared to those samples obtained from access tissues in patients with brain tumors. However, the long-term sustainability was slightly higher in access tissue from patients with brain tumors making those cases excellent candidates for long-term cultivation of brain tissue after proper and critical selection.

### Possible applications to study the physiology of the human brain

Because of the isolation of the cells from the brain and the exposure to artificial culturing settings, it should be noted that human organotypic brain slice cultures are most likely undergoing several changes once they are in culture. Nevertheless, we and others have shown a remarkable stability of fundamental properties of the neurons over time [23], [36], [37]. The extended experimental time period opens up the possibility to use genetic tools for manipulation such as optogenetics [37], [39] and chemogenetics [37] and use experimental paradigms involving longer time frames such as plasticity responses or gene expression alterations.

### Possible applications for pathological models

The described protocol for long-term human organotypic brain slice cultures provides a comprehensive framework to study not only physiological neuronal activity, but can be also used to investigate different pathological conditions. Through genetic modulation of ion channels or insertion of a lesion within the cultured slices, one can obtain various disease models mimicking epilepsy or neurogenerative diseases [40], [41]. Even implantation of tumor cells (i.e. glioblastoma) into the human brain slices can be successfully performed in order to create a tumor model that resembles the natural human brain microenvironment [42], allowing the investigation of both tumor growth patterns as well as treatment regimes.

## Conclusion

Neuronal viability within organotypic human brain slice cultures can be maintained for several days up to 3 weeks providing a unique experimental environment in the field of neuroscience. Although a good interdisciplinary set-up is an important prerequisite for the successful establishment of human brain slice cultures, the described method can be easily adapted and exploited in a laboratory setting, opening a novel and powerful experimental neuroscientific framework.

## Acknowledgements

This work was supported by the Confocal Microscopy Facility, a core facility of the Interdisciplinary Center for Clinical Research (IZKF) Aachen within the Faculty of Medicine at RWTH Aachen University.

## CRediT author statement

**Aniella Bak:** Conceptualization, Methodology, Validation, Data Curation, Formal analysis, Investigation, Writing – Original Draft, Review & Editing, Visualization; **Henner Koch:** Conceptualization, Methodology, Software, Validation, Formal analysis, Writing – Original Draft, Review & Editing, Resources, Supervision, Project administration, Funding acquisition; **Karen M. J. van Loo**: Methodology, Validation, Writing – Review & Editing, Supervision; **Katharina Schmied**: Methodology, Validation, Investigation, Data Curation; **Birgit Gittel**: Investigation, Validation; **Yvonne Weber**: Supervision, Project Administration, Funding acquisition; **Niklas Schwarz:** Methodology, Writing – Original Draft; **Simone Tauber:** Resources, Writing – Review & Editing; **Thomas V. Wuttke:** Conceptualization, Methodology, Software, Validation, Formal analysis, Writing – Original Draft, Review & Editing, Resources, Supervision, Project administration, Funding acquisition; **Daniel Delev:** Conceptualization, Methodology, Software, Validation, Resources, Writing – Original Draft, Review & Editing, Supervision, Project administration

## Supplement A: Notes

### Note 1: Slicing solution alternatives

We use a slicing solution (s-aCSF) that was adopted from protocols described for acute human slices [29]. Nevertheless, other slicing solutions have been reported and might be a feasible alternative [43].

### Note 2: Sterility & ethanol

Sterility is key when preparing brain slice cultures. Hence, every tool or object used for slice preparation should be sterilized beforehand, either by UV light, heat or ethanol. This includes objects such as absorbent linings, slicing chamber, preparation dishes, pipette tips, glue, tools to operate the vibratome and every other object listed in the protocols. Objects that are difficult to sterilize, such as nylon nets or bubble stones, should be exchanged for more practical options (e.g. cannulas instead of bubble stones).

Whenever ethanol is used for sterilization, it should not be brought into contact with the slicing solution or the tissue itself. Furthermore, anything disinfected with ethanol should be left for proper evaporation before usage.

### Note 3: hCSF precautions

- **Collection & Processing**: hCSF should be processed immediately after extraction from the patient. After processing, hCSF can be stored at -80°C for several months. Discoloration of hCSF at the time of collection is sometimes possible due to bleeding or high protein content and not to be seen as a quality indicator, as long as osmolarity is still in range. Nevertheless, hCSF should always be clear and never appear cloudy.
- **Infection risk**: One should always be aware that hCSF and human tissue samples are potentially infectious and could harbor life-threatening pathogens such as HIV or Hepatitis virus. In consequence, hCSF as well as brain tissue should always be handled with great care and safety measures (gloves, avoiding contact, do not handle with open wounds) accordingly.
- **Batch segregation**: Larger volume batches from individual patients should be made distinguishable from each other by keeping them segregated and labeling them accordingly. In case of contamination or ineptness, the affected batch can thus be excluded from culturing processes. Smaller volume batches of hCSF can be pooled together to level out any elevated or reduced parameters and create an even batch of hCSF to avoid exposition of the brain slice cultures to permanent osmotic and component level changes.

### Note 4: Hypothermia and re-warming

Human brain tissue should be kept at hypothermic conditions from the time point of resection until the tissue has been sliced to help slow down degrading processes and support slice stability. As soon as the slicing of the tissue is finished, an optional gentle rewarming can be performed by keeping the slices in room temperature solution for additional cutting steps (downsizing and removal of damaged/incomplete parts) and subsequently transferring them to a slice keeper with pre-warmed aCSF (30 – 33°C) until culturing. Brain slices intended for direct acute experiments or recordings should be left for recovery and rewarming in aCSF at 30 - 33°C for at least 1h.

### Note 5: Oxygenation

Constant carbogenation/oxygenation is vital for the survival of cortical brain tissue. From the time point of resection until the brain slices are placed on the membrane inserts for culturing, the brain tissue needs to be constantly submerged in a carbogenated solution. If there are working steps, such as the transfer to the culture hoods, which do not allow carbogenation, the time frame should be kept to an absolute minimum to avoid hypoxia and therefore neuronal death.

### Note 6: Choice of suitable surgeries

The choice of the proper surgery builds the basis for the further success of the experiments. Normally, one should try to obtain the part of the tissue used for surgical approach which appears to be as physiological as possible. This will guarantee higher success rates in brain tissue culturing given the uncompromised cortical structure. Therefore, tissue acquired from temporal lobe resection either due to mesial temporal lobe epilepsy or to Meningioma or Cavernoma lesions that are usually well-circumscribed and if the perilesional tissue is not compromised by swelling or hemosiderin provide the best tissue for organotypic brain cultures. Tissue derived from malignant lesions like glioma and metastasis may be compromised by tumor infiltration or pronounced edema. If such tissue can be obtained, then it should be as far as possible from the malignant lesion.

### Note 7: Surgical considerations

The most important surgical considerations can be summarized withing the following three aspects:

a. Do not coagulate the cortex!
b. Avoid tissue manipulation.
c. Reduce the time between tissue disconnection from the rest of the cortex and its transfer into the medium.

### Note 8: Block/Slice handling

The correct handling of slices is an absolutely crucial step to obtain viable and intact brain slice cultures. Brain slices are very delicate and should always be handled with great care.

- Brain tissue blocks as well as slices should never be squished or exposed to mechanical stress of any kind.
- In order to cut brain tissue blocks or slices, clean single cuts with sharp tools should be utilized.
- When mechanically manipulating brain tissue blocks or slices, any sharp tools or objects should be avoided, as they will damage the tissue immediately. Brain tissue should always be moved or transferred with blunt tools only in order to avoid damaging or tearing.
- Brain tissue should always be fully submerged in liquids while processing, as it ensures the integrity of the blocks and slices by avoiding mechanical stress and damage to the tissue.
- Brain slices should not be exposed to excessive air blowing, e.g. by the carbogenation system or trying to remove them from the block while slicing. Please behold to the Notes section ‘Slicing difficulties’.
- During slicing, the tissue should only be moved through the different dissection dishes and bottles “one-way”. Tissue should never be placed in its previous dish to avoid contamination and exposure to cell debris and other damaging agents within the slicing solutions.

### Note 9: Slicing difficulties

During slicing of the brain tissue, several difficulties might occur.

- **Orientation**: Slicing should be performed perpendicular to the cortical surface and parallel to cortical gyri to obtain slices spanning all cortical layers. If cortical layer 4 (presenting as a lighter line across the slice) or the white matter (presenting clearly delimited from the gray matter) appear to be blurred, the orientation should be re-adjusted. Alignment of blood vessels with sectioning plain perpendicular to the cortical surface can also serve as an indicator of correct orientation.
- **Pushing**: In some cases, the brain slice can get attached to the razor blade, resulting in the blade ‘pushing’ the slice rather than cleanly slicing through it, leading to uneven thickness or tearing of the slices. This occurrence often correlates with pieces of the Pia mater still being attached to the brain tissue block and, as a result, clinging to the razor blade. Once this phenomenon occurs, it can only be resolved by reducing the slicing speed and sometimes carefully cutting or peeling off remaining pieces of pia mater with microdissection forceps of microscissors. If this does not solve the problem and brain slices are still cut unevenly sliced, exchanging the tissue block is recommended. The problem may also occur in case the block started to detach from the agar surface, if superglue was spread to the sides of the block or once the bottom of the block and the gluing surface are approached.
- **Detaching slices**: Usually, brain slices should detach from the tissue block automatically once the razor blade sliced through the tissue block completely (up to or even slightly into the agar cube mounted behind the tissue block) and slices should not be detached before the sweep is fully completed. In cases in which the brain slice does not detach, the razor blade should be stopped while still covering the tissue block and before reverting to its starting position. In this way, the remaining tissue is protected from tools used to detach the brain slice.
- **Slice integrity**: Every brain slice should encompass all cortical layers. If a slice is damaged at the cortical layer or elsewhere, the compromised parts should be removed with a clean cut to keep the tissue viable.

### Note 10: Infections

If the slicing process is performed on a working bench outside a sterile hood, fungal and bacterial infections of the cultured tissue can be a prominent problem. As the hCSF used for culturing does not contain any antibiotic/antimyotic agents, precautions should be taken to avoid infections.

- The working space should be cleaned thoroughly beforehand with 70% ethanol and every single object/tool used for the slicing process should either be sterilized by UV, heat or ethanol.
- The slicing process should always be performed as if it took place under a sterile hood, meaning the performing persons should wear gloves and sanitize them before starting or continuing work at the working space. Surgical mask, hair net and surgery clothing can be of advantage as well to avoid possible contaminations.
- The number of persons present at the working space during the slicing process should be kept to a minimum. Doors should be closed whenever possible.
- The slicing medium should contain at least antibiotics, but preferably both antibiotic and antimyotic agents to prevent infections.
- Tubing and cannulas for the carbogenation system should always be new and sterile.
- If needed, antibiotic and/or antimyotic agents can be added to the hCSF for the first culturing step and replaced by pure hCSF during the next change of medium.
- Silicone preparation dishes should be exchanged regularly, depending on use frequency, as an increasing amount of cuts in the silicone can increase the risk of infections.

### Note 11: Qualitative evaluation of cultures

The macroscopical evaluation of the viability of human brain slice cultures poses some difficulties in comparison to rodent cultures, as they will only thin out, shrink or change their appearance marginally during degradation, at least during the first few days. Nevertheless, there are some indicators of slice culture viability that can be assessed macroscopically.

- **Membrane adherence**: Viable brain slices will adhere to the semipermeable membrane of the culturing insert during the first 1-2 days in culture. This adherence is not as robust as seen in rodent brain cultures, as human brain slices are still easily removable from the membrane with a soft brush, however, brain slices should not float or detach immediately if viable. The goal of this culturing method is to position the brain slice at an air-liquid-interface, meaning the slice should always be humid on top of the insert, but not covered in fluid to maintain air contact.
- **Appearance**: Human brain slices usually exhibit a clear shine on their surfaces if the culturing is successful. If the slices appear dull, it can be due to possible degradation or insufficient humidity levels within the incubator.
- **hCSF**: The hCSF used as culturing medium should always appear clear. If the medium appears cloudy, it is either due to cell debris and therefore tissue degradation, or any kind of infection. Tissue degradation as well as infections can also manifest as residue on top of the culturing insert. Please note that a mild degree of clouding can be normal directly after slicing as the slices could be still discharging cell debris originating from the cutting borders, but should not reoccur after the first medium change.

Additionally, quality and viability of cultures can be either assessed by electrophysiological recordings on single-cell or network level, or by inspection of fluorescent expression after viral transduction.

### Note 12: Viral transduction

- The application of the virus is a potential source of contamination. Therefore, work here should also be as clean as possible and the opening of the culture plates should be reduced to a minimum.
- Surface application of the virus should be performed carefully without touching or even penetrating the slice with the pipette tip. There should be no air bubbles on the top of the slice after applying the virus droplet.
- For easier pipetting, the virus can be diluted 1:1 or 1:2 with PBS. Nevertheless, application of > 5μl should be avoided to allow the droplet to fully remain on top of the slice instead of overflowing and dispersing on the sides.
- Viral transduction of one slice with multiple, different viruses is possible.
- Transduction within the first 24 - 48 h after the start of culturing is recommended, as this allows optimal use of the subsequent time for expression of the transduced payload and planned experiments. Additionally, it allows the virus to penetrate the tissue before formation of a glial scar, which could impair transduction efficacy.

